# Weathering the storm for love: Mate searching behaviour of wild males of the Sydney funnel-web spider (*Atrax robustus*)

**DOI:** 10.1101/2024.07.23.604707

**Authors:** Caitlin N. Creak, Hugo Muirhead, Michael Kasumovic, Russell Bonduriansky, Bruno A. Buzatto

## Abstract

The risky business of mate-searching often leaves the actively searching sex facing threats and rapidly changing conditions. Yet, active mate-searching behaviour is rarely studied in invertebrates, and we have limited understanding of how mate-searching strategies have evolved to cope with risks posed by harsh weather. We investigated how mate-searching males move through their habitat and how their movement is affected by weather conditions in the Sydney funnel-web spider (*Atrax robustus*), one of the world’s most venomous spiders. As is common in mygalomorphs spiders, females are functionally sessile, and are thought to spend their whole lives in a single burrow, whereas males must permanently abandon their burrows to mate during the breeding season. Nineteen male spiders were fitted with micro-radio transmitters and tracked during their mating seasons in 2020 (n = 2), 2021 (n = 8) and 2022 (n = 9) in Lane Cove National Park, in Sydney, Australia. Males moved at night, typically in a zig-zag pattern, and were found in new locations on approximately 50% of daily resighting’s. Males often spent several days in a female’s burrow, and some female burrows were visited by multiple males. When outside a female’s burrow, males constructed and occupied temporary shelters (‘temporacula’). Males were most likely to move and/or moved furthest when there was no rain, and on warm nights after cool days. Our findings suggest that mate-searching *A. robustus* males prefer to search for females in less risky conditions, revealing novel risk-minimizing strategies, especially in response to rainfall and temperature.

## Introduction

Mate searching is one of the most dangerous behaviours for many species (Kasumovic et al., 2007). Numerous types of mate-searching strategies are found across different taxa and environments, including visual searching (Hemptinne et al., 1996), targeting host plants or animals (Estrada & Gilbert, 2010; Mulrey et al., 2015), following pheromonal trails (Charlton & Card, 1990) and responding to auditory or vibrational signals (Cocroft & Rodríguez, 2005). Many animals use combinations of mate searching strategies, as such a multimodal approach to finding a mate is assumed to increase individual mate searching success (Rypstra et al., 2009). Multimodal mate searching is believed to be especially advantageous in complex or changing habitats because it allows for facultative behavioural adjustment of strategies depending on specific conditions (Brandt et al., 2018). Understanding how environmental factors shape the evolution and expression of mate-search strategies could also help predict how anthropogenic changes can affect these strategies (Heuschele et al., 2012).

The evolution of mate searching strategies is expected to optimise the trade-off between the cost of searching and the probability of success (Sullivan-beckers & Hebets, 2014). Higher risk and energy expending behaviours may be costly for the survival of individuals searching for a mate but will still be favoured if the success in finding receptive high-quality mates is greatly enhanced. Despite the available knowledge on sexual selection and mate searching strategies in general (Fromhage et al., 2016; Parker, 1978), many questions remain about how mate searching strategies are shaped by the environment and how such strategies vary among animals across diverse life histories and ecological niches. One aspect of mate-searching in particular – the active searching stage before locating a mate – remains little-studied, and the strategies and costs involved poorly understood.

Mate searching in spiders is of particular interest due to the role that their ecological niche is expected to play in how mate searching strategies have evolved. In many species of araneomorph spider, aerial webs are created in complex vegetation or habitat (Dimitrov & Hormiga, 2021). These webs often serve multiple purposes, such as prey capture, predator warning, and shelter, and are also used in courtship and mating signals (Copperi et al., 2019; Dimitrov & Hormiga, 2021). In web-dwelling species, sexually mature males must abandon their own web in order to search for a conspecific female’s web in a complex, 3-dimensional environment (Kasumovic & Andrade, 2004). For example, mature female western black widow spiders *Latrodectus hesperus* are widely distributed and do not leave their webs unless provoked, yet the much smaller mature males abandon their webs to search for mates and are able to locate and distinguish between virgin and mated females by following airborne pheromones emitted by females. Once in reach of a female’s web, males also use contact pheromones to assess female mating status (Kasumovic & Andrade, 2004).

Despite the considerable amount of research on mate searching and communication across different species of spiders (Herberstein et al., 2014), few studies have explored mate searching and the traits associated with success in burrowing (fossorial) spiders. It remains unclear what strategies are typically employed by mate-searching individuals in burrowing spiders, or how such strategies compare with other groups of spiders. Several studies have investigated the courtship and mating behaviours of wolf spiders (Araneomorphae : Lycosidae), a mostly fossorial group (Carballo et al., 2017; Carrel, 2003; Marshall, 1995). More recently, the courtship and mating behaviour of the Sydney funnel-web spider *Atrax robustus* (Mygalomorphae: Atracidae) were also described in detail (Frank et al. 2023). However, nothing is known about the searching strategies of actively mate searching Lycosids or Atracids in the wild. Exploring mate searching and communication in an ancient group of spiders such as mygalomorphs offers a valuable opportunity. Many mygalomorph species are relatively large and robust, allowing individual tagging and tracking (Janowski-bell & Horner, 1999). Understanding the variation in mate searching across the spider phylogeny will also provide insight into the evolution of the traits associated with successful mate searching in different groups of spiders.

The majority of mygalomorph spiders dig burrows in the ground, in which they spend most of their lives, up to 42 years for the females of some species (Mason et al., 2018). Burrows differ greatly in morphology between genera and species, but are typically lined with silk (Wilson et al., 2023). Some species of mygalomorph create trapdoors, while others may have multiple entrances, dead ends or other features (Wilson et al., 2023). Burrows provide shelter from predators and weather, retain moisture, and create ambush positions for prey capture (Cloudsley-Thompson, 1983; M’rabet et al., 2007). During their mating season, male mygalomorphs abandon their burrows permanently in search of mates, spending one season mating and then perishing (Mason et al., 2018). During this period of active mate searching, male behaviour has only been documented in two species of mygalomorph. One study tracked 11 males of the tarantula *Aphonopelma hentzi* (Theraphosidae) for between three and 24 days during their mating season, and showed that males moved up to 1.3 Km, in an apparently random pattern with no discernible direction (Janowski-Bell & Horner, 1999). In two separate studies, male *Aphonopelma anax* were tracked with the aim of describing male thermoregulation and metabolic rate during their mating season. In 2002, 14 males were tracked every two hours for two consecutive days (Shillington, 2002). In 2004 & 2005, 26 males were tracked between one and 34 days (Stoltey & Shillington, 2009). During these studies, males of *A. anax* were observed moving up to 365 meters per day, and it was found that males successfully thermoregulated with the use of temporary day time retreats, so that their resting metabolic rate remained constant over the duration of the mating season.

Our goal in this study was to better understand mate searching behaviour in the Sydney funnel-web spider *Atrax robustus* (Atracidae)(Gray, 2010), a ground-dwelling mygalomorph endemic to south-eastern Australia. *A. robustus* has a small species range located along the coast of the Greater Sydney metropolitan area (NSW), and can be classified as a short-range endemic (SRE), occupying an area of less than 10,000 km² (Gray, 2010; Harvey, 2002). A visually intimidating species, mature males are glossy black, have the cephalothorax and legs relatively glabrous, and a densely haired abdomen (Gray, 2010). Females tend to have a similar appearance to males but are generally larger, and their abdomen varies in colouration from black to brown (Gray, 2010). After moulting into the adult stage, at approximately 5 – 7 years of age (Braxton Jones, pers communication), *A. robustus* males permanently leave their burrow in search of females during the warmer months of the year (Bradley, 1993).

During the mate-searching period, male *A. robustus* come into frequent contact with humans (Isbister & Fan, 2011). Sexually mature male *A. robustus* are arguably the most venomous species of spider currently known, possessing a neurotoxic venom containing peptides called δ-hexatoxins (δ-HXTXs-Ar1a) (Herzig et al., 2020). Although some animals survive envenomation (Nicholson & Graudins, 2002), humans and other primates are particularly susceptible. In humans, envenomation symptoms begin with numbness and quickly progress to hypertension, pulmonary edema and ultimately death (Isbister et al., 2005). While this venom has a potentially lethal effect on humans, there have been no recorded human deaths since the introduction of an antivenom in 1984 (Hartman & Sutherland, 1984). Females and juvenile *A. robustus* pose less risk to humans, as their venom contains lesser quantities of δ-HXTXs-Ar1a (Isbister et al., 2005). The toxicity of male *A. robustus* venom towards humans highlights the importance of understanding the behaviour of mature males during the mating season. Understanding their movement patterns can reduce the likelihood of contact between *A. robustus* and humans, and perhaps reduce the fear associated with these spiders.

To understand the mate searching behaviour in male *A. robustus*, we investigated how these males move through their environment during their mating season. Our aims were first to describe male long-distance movement by examining distances travelled, time spent in burrows, changes in movement over time, and the plant community types traversed. Because males are likely to search for females at long distances using volatile pheromones (Scott et al., 2018), we predicted that male movement would be generally linear at a larger scale and exhibit a zig-zag pattern at a smaller scale, as males follow pheromone plumes towards female burrows. Our second aim was to investigate the effect of environmental variables, such as precipitation and temperature, on male movement. Theory predicts that mate-searching males will be more likely to move, or move longer distances, when the costs of moving are low, such as when weather conditions are optimal. It is often assumed that rainfall encourages greater *A. robustus* activity, but this could be due to the findings of Bradley (1993), who reported that after heavy rainfall an increased number of female *A. robustus* were collected from pitfall traps, presumably a result of those females being dislodged from their flooded burrows, since females normally do not abandon their burrows. However, that study did not assess the activity of males after heavy rainfall. In the event of heavy rain, males outside of a burrow are likely under higher risk of injury or death by flood water, debris and potentially even predators, it is therefore possible that males may avoid mate searching in risky conditions.

## Methods

### Collection of wild spiders

This study was conducted on Eora and Kuring-gai (also Guringai) Nations’ lands. Individual male *Atrax robustus* specimens (n = 19) were collected between February and April of 2020 (n = 2), 2021 (n = 8), and 2022 (n = 9), from Lane Cove National Park (−33.761, 151.106), Sydney, Australia. To collect males, trips on foot after 8pm AEST were made along multiple fire trails within the national park. Males were spotted on and alongside the fire trail in undergrowth and upon elevated roadside banks. Collecting individuals involved placing an open 500ml plastic container over the top of the spider, then encouraging the spider to walk into the container using the lid. Once collected, we labelled the location using flagging tape and saved the GPS location of each individual using a Garmin GPSMAP 62sc.

### Transmitter attachment

We used model T15 micro-radio transmitters (Advanced Telemetry Systems, Gold Coast, Qld., Australia) to tag spiders (Figure 1). The T15 transmitters weigh 0.15 grams and are 3.4mm x 11mm for the main body (Figure 1 A) plus a 150mm flexible antenna. Transmitters had a battery life of approximately 30 days. Two ATS R410 scanning receivers with a 3-element folding yagi directional antenna were used to locate individuals with transmitters.

**Figure 1:**
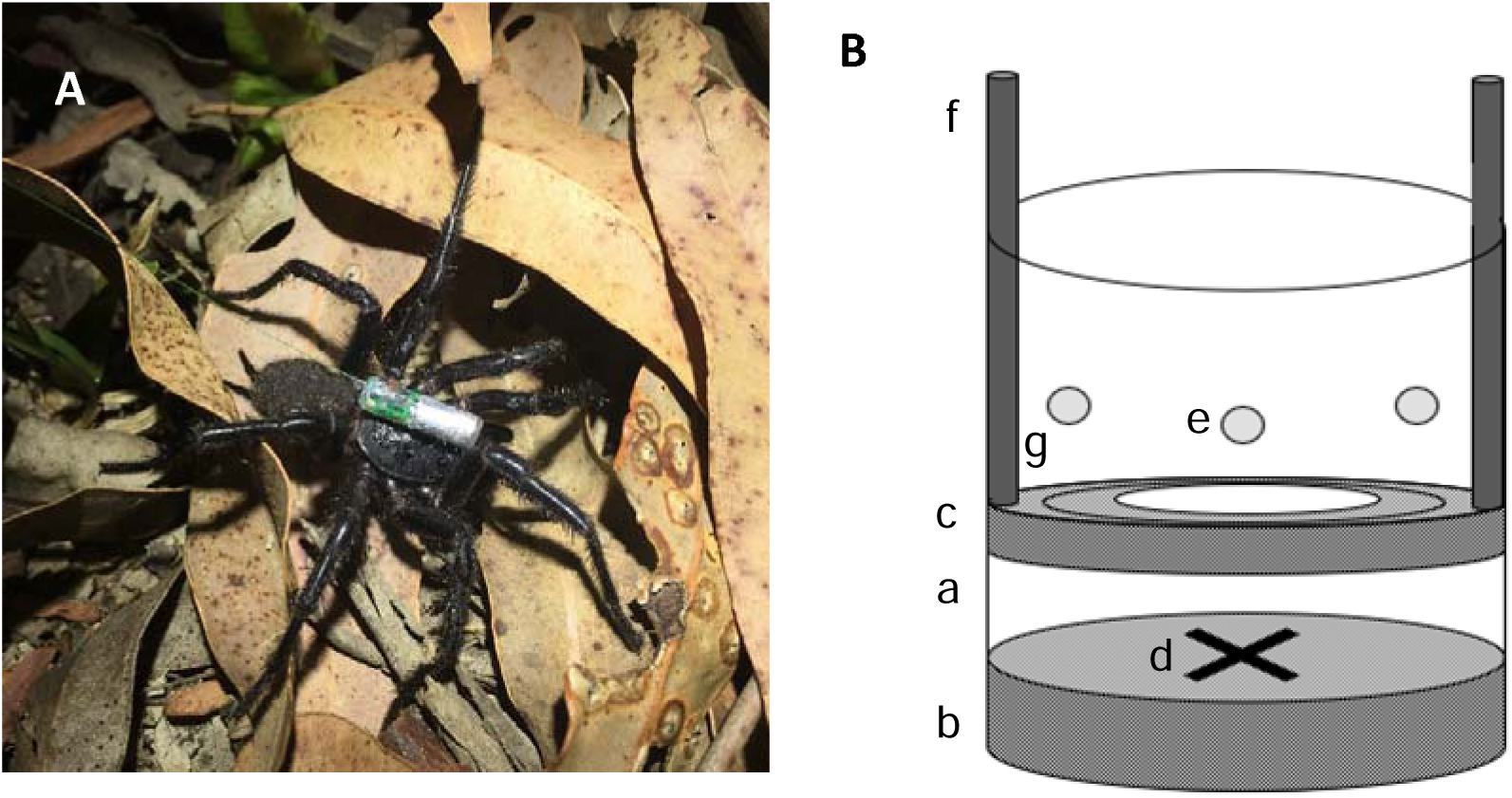
A) Photograph of a male with a T15 transmitter attached to the cephalothorax. Taken by Bruno Buzatto. B) Diagram of the apparatus built for attaching the transmitters; a; 500 ml cylindrical container. b; base sponge. c; upper sponge. d; cross marked on sponge to indicate spider cephalothorax placement. e; air holes. f; dowel rods. g; persepex plate.

At first capture, individuals were taken back to the lab at UNSW Sydney and photographed inside their container. In 2022, males were also weighed using a HT-120 compact balance (A&D Ltd., Thebarton, SA, Australia). Individuals were given a minimum of 12 hours to settle and were provided with water ad libitum before the transmitter was attached. Each ATS T15 transmitter was switched on using the ATS T15 controller app immediately prior to attachment to avoid battery life wastage. To attach the transmitters, each individual spider was placed into a 500ml plastic container with holes drilled into the sides and lid (Figure 1 B). A damp sponge was placed in the bottom of the container to ensure individuals did not desiccate. Once the individual was transferred to the container, the container was placed into a sealed plastic bag, which was then filled with CO_2_ gas and sealed off for three minutes or until the individual was lethargic and easily manoeuvrable.

The individual was them moved using forceps to the centre of the sponge (location marked with an X in Figure 1 B), and a second sponge (cut into a ring shape with a Perspex plate around the outside and two dowel rods attached to it) was placed over the top of the individual to expose the cephalothorax but restrain the legs and abdomen (Figure 1 B). A blunt size 28 tapestry needle was then used to smear a small amount of 430 Loctite super glue (Henkel Corp., Ohio, USA) onto the cephalothorax and then onto the transmitter itself. Using fine point forceps, the transmitter was placed onto the cephalothorax and held there for approximately 15 seconds until the glue set. The cephalothorax was chosen as the site of transmitter attachment because it is a non-porous sclerite, whereas the abdomen is a membranous structure that expands and retracts and thus attempting to attach the transmitter on it would injure the spider.

The top sponge was then removed, and the spider remained in the container for 30 minutes to allow the glue to cure further. The spider was then moved into its original container and allowed to recover for a minimum of 12 hours. Individuals were released in the location where they were found 12 – 48 hours after the transmitter was attached in the lab. In three cases, individuals were kept in the lab for seven days as they were collected immediately prior to the field site flooding and to ensure a safe release and safe field practices, we allowed extra time after the flood had passed.

### Tracking of tagged spiders in the wild

Individuals were tracked one at a time every day using the receiver and antenna, unless extreme weather events prevented entry into the National Park. We recorded details of each individual on every day that they were tracked once the location of an individual was narrowed down to < 1 square meter or the individual was sighted. In each case, we recorded a GPS coordinate and secured labelled flagging tape to foliage where the spider was located. We next measured the distance between the previous and current location (“nightly distance moved”) using a retractable 60 m measuring tape. If individuals were tracked to the same location and had not moved, this was recorded as a nightly distance moved of 0 m. If an individual had not moved and had not been physically sighted after three days, the site was thoroughly examined to ensure the transmitter had not dislodged or that the individual had not died. All males were tracked until either their transmitters dislodged (n=15), the transmitter ran out of battery (n=2), or the spider was found dead (n=2). The number of days of tracking and number of observations for each individual spider are shown in Table 1. If males were tracked into a burrow, the GPS coordinates of the burrow were recorded, and the individuals were left undisturbed. All males that were tracked into burrows subsequently re-emerged from the burrows.

**Table 1:**
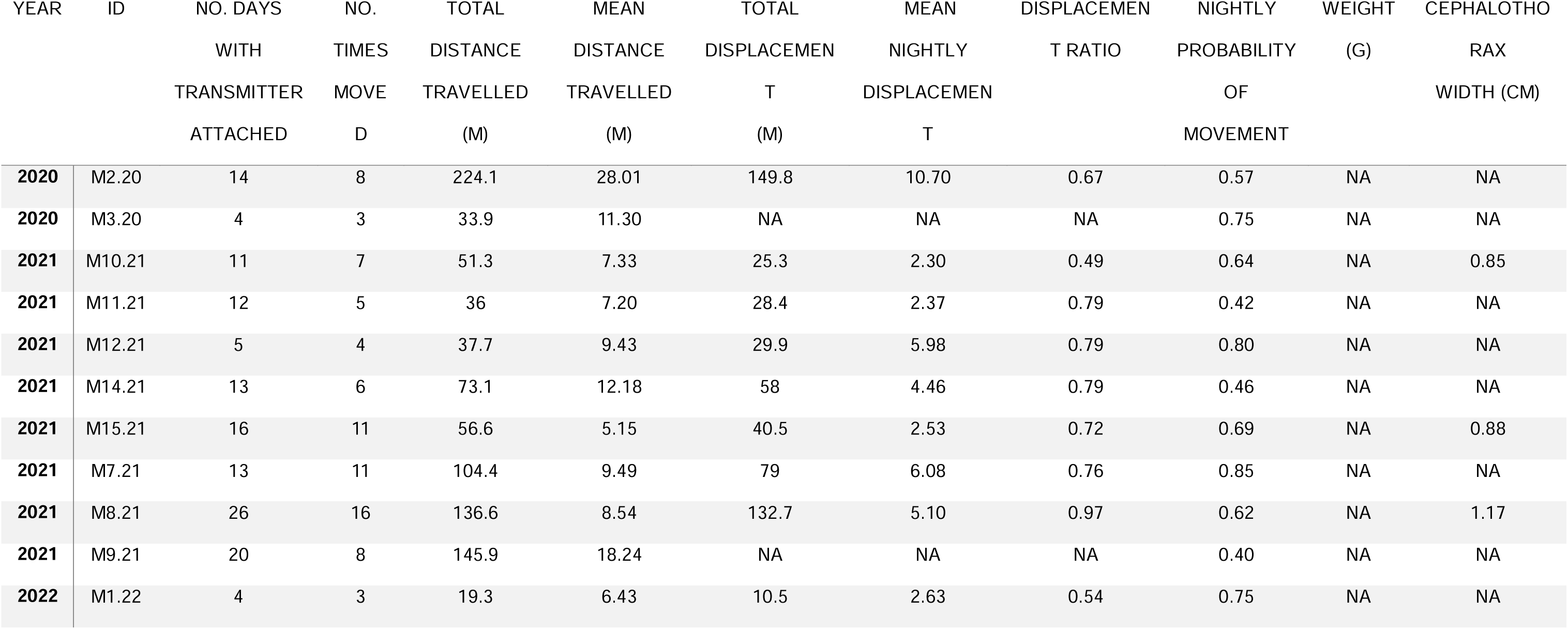
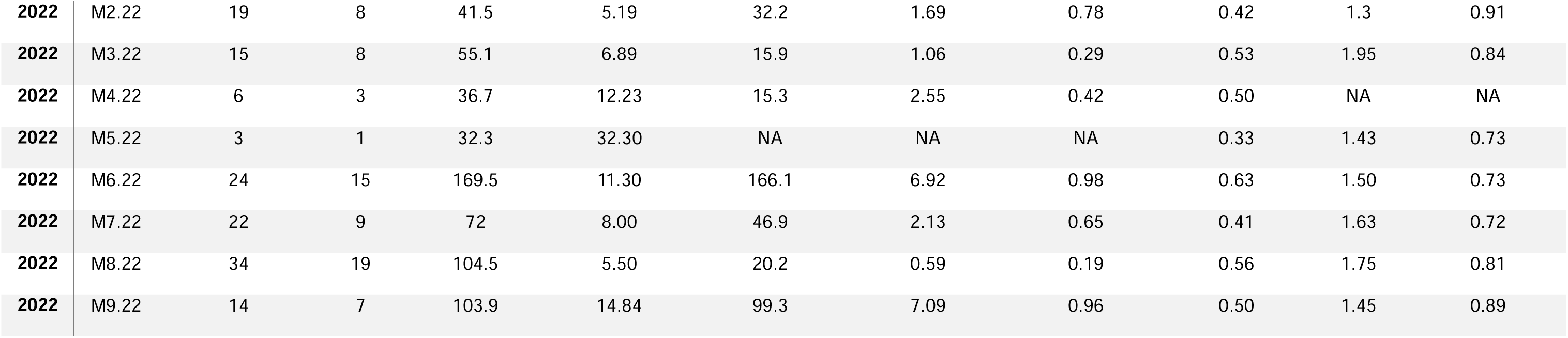
Summary of movement data for individual spiders that were tracked over the three years of this study: **Year** = year individuals were tracked, **ID** = unique identification number for each individual spider, **No. of observations** - the total number of times each individual had a new nightly distance recorded. **No. of Days with transmitter attached** = total number of days that each individual carried a transmitter, beginning at release and ending with transmitter dislodge, battery expiry or death of the individual, **Total distance travelled** = the total number of meters moved recorded for each individual, **Mean distance travelled** = the total distance divided by the number of days the individual moved, **Total displacement** = distance between first and last GPS location (this could not be measured for three individuals due to GPS error), **Mean nightly displacement** = Total displacement divided by the number of days with transmitter attached, **Displacement ratio =** Displacement divided by total distance travelled, **Probability of movement** = number of days when the spider moved divided by the total number of days with the transmitter attached, **Individual weight** = weight at time of initial capture (mg), **Cephalothorax width** = measurement of the widest part of the cephalothorax, between legs II &III (cm).

| YEAR | ID | NO. DAYS<br>WITH<br>TRANSMITTER<br>ATTACHED | NO.<br>TIMES<br>MOVE<br>D | TOTAL<br>DISTANCE<br>TRAVELLED<br>(M) | MEAN<br>DISTANCE<br>TRAVELLED<br>(M) | TOTAL<br>DISPLACEMENT<br>T<br>(M) | MEAN<br>NIGHTLY<br>DISPLACEMENT<br>T | DISPLACEMENT<br>RATIO | NIGHTLY<br>PROBABILITY<br>OF<br>MOVEMENT | WEIGHT<br>(G) | CEPHALOTHORAX<br>WIDTH (CM) |
| --- | --- | --- | --- | --- | --- | --- | --- | --- | --- | --- | --- |
| 2020 | M2.20 | 14 | 8 | 224.1 | 28.01 | 149.8 | 10.70 | 0.67 | 0.57 | NA | NA |
| 2020 | M3.20 | 4 | 3 | 33.9 | 11.30 | NA | NA | NA | 0.75 | NA | NA |
| 2021 | M10.21 | 11 | 7 | 51.3 | 7.33 | 25.3 | 2.30 | 0.49 | 0.64 | NA | 0.85 |
| 2021 | M11.21 | 12 | 5 | 36 | 7.20 | 28.4 | 2.37 | 0.79 | 0.42 | NA | NA |
| 2021 | M12.21 | 5 | 4 | 37.7 | 9.43 | 29.9 | 5.98 | 0.79 | 0.80 | NA | NA |
| 2021 | M14.21 | 13 | 6 | 73.1 | 12.18 | 58 | 4.46 | 0.79 | 0.46 | NA | NA |
| 2021 | M15.21 | 16 | 11 | 56.6 | 5.15 | 40.5 | 2.53 | 0.72 | 0.69 | NA | 0.88 |
| 2021 | M7.21 | 13 | 11 | 104.4 | 9.49 | 79 | 6.08 | 0.76 | 0.85 | NA | NA |
| 2021 | M8.21 | 26 | 16 | 136.6 | 8.54 | 132.7 | 5.10 | 0.97 | 0.62 | NA | 1.17 |
| 2021 | M9.21 | 20 | 8 | 145.9 | 18.24 | NA | NA | NA | 0.40 | NA | NA |
| 2022 | M1.22 | 4 | 3 | 19.3 | 6.43 | 10.5 | 2.63 | 0.54 | 0.75 | NA | NA |
| **2022** | M2.22 | 19 | 8 | 41.5 | 5.19 | 32.2 | 1.69 | 0.78 | 0.42 | 1.3 | 0.91 |
| **2022** | M3.22 | 15 | 8 | 55.1 | 6.89 | 15.9 | 1.06 | 0.29 | 0.53 | 1.95 | 0.84 |
| **2022** | M4.22 | 6 | 3 | 36.7 | 12.23 | 15.3 | 2.55 | 0.42 | 0.50 | NA | NA |
| **2022** | M5.22 | 3 | 1 | 32.3 | 32.30 | NA | NA | NA | 0.33 | 1.43 | 0.73 |
| **2022** | M6.22 | 24 | 15 | 169.5 | 11.30 | 166.1 | 6.92 | 0.98 | 0.63 | 1.50 | 0.73 |
| **2022** | M7.22 | 22 | 9 | 72 | 8.00 | 46.9 | 2.13 | 0.65 | 0.41 | 1.63 | 0.72 |
| **2022** | M8.22 | 34 | 19 | 104.5 | 5.50 | 20.2 | 0.59 | 0.19 | 0.56 | 1.75 | 0.81 |
| **2022** | M9.22 | 14 | 7 | 103.9 | 14.84 | 99.3 | 7.09 | 0.96 | 0.50 | 1.45 | 0.89 |

### Ethical Note

No ethics permit was required for this study because spiders are not covered by animal ethics legislation in Australia. However, the methodology designed for this study aimed minimize potential harm to the spiders. No licence was required by National Parks as nothing was being permanently removed or altered in the field site.

During CO_2_ exposure, the spiders did not elicit any signs of discomfort (as discussed by Dombrowski et al., 2013). Each spider recovered normally in a dimly lit, quiet environment, and subsequent behaviour during the recovery period appeared normal. Only one fatality occurred using this procedure due to excess glue being applied. This individual was humanely euthanised using CO_2_. The transmitters were an average of 9% of spider body weight, ranging from 7.7% - 11.5%. Most of the tagged spiders lost their transmitters while still alive. In the two cases where the transmitters ran out of battery while attached to a spider, both males had already been monitored for >25 days and exhibited normal behaviour throughout the study. Rather than attempt removing the transmitter, potentially disturbing natural behaviour or harming these individuals, it was deemed safer to leave the trackers attached as the males of this species die at the end of their mating season regardless.

### Statistical analysis

All statistical analyses were conducted using R version 4.3.1 (2023). Information on plant community types was obtained from the NSW State Vegetation Type Map (State Government of NSW and NSW Department of Climate Change, Energy, the Environment and Water, 2022) then layered over maps (configured in QGIS 3.30.3) containing data on GPS locations for individual spiders. Temperature and rainfall data were obtained from The Australian Government, Bureau of Meteorology (Australian Government, Bureau of Meteorology, 2023). The data consist of minimum and maximum temperature and rainfall in all forms of precipitation recorded at 9am AEST, and therefore represent weather conditions for the preceding 24 hours.

Total distance travelled over the course of the season was calculated for each spider as the sum of its nightly movement distances, and the mean nightly movement distance was calculated by dividing the total distance by the number of nights when the spider moved to a new location. In addition, we estimated the linear projection (“displacement”) of each spider’s movement over the course of the season by measuring the distance between the first location of capture and the last location recorded in QGIS. To calculate how far on average each male moved per night away from the initial place of capture, we divided displacement by the number of nights between the date of release and the date of final observation (“mean nightly displacement”, Table 3). To quantify the linearity of movement, we then divided displacement by the total distance travelled (“displacement ratio”, Figure 2, Table 3). The displacement ratio would equal 1 if the spider moved in a straight line away from its location of capture, and 0 if the spider circled back around to its initial capture location.

**Figure 2.**
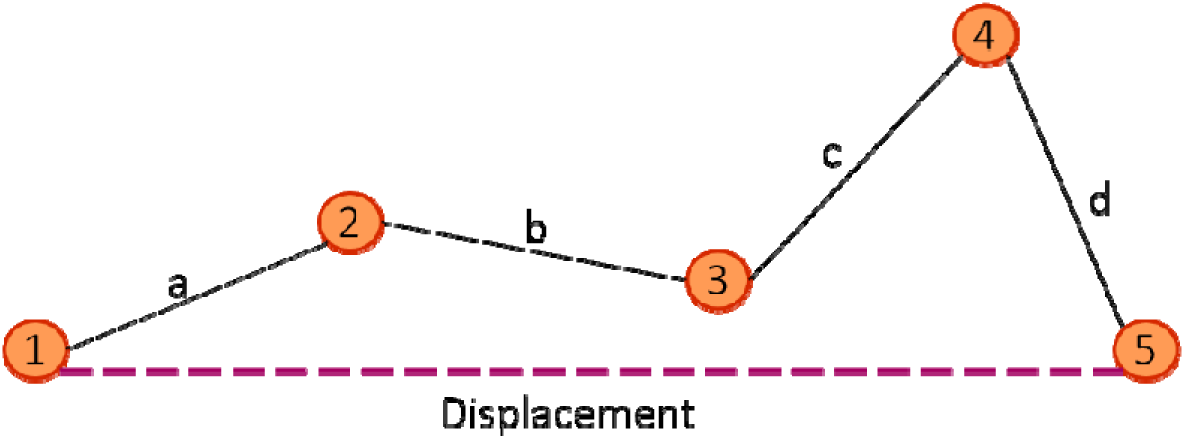
Visual representation of how movement metrics were collected: distances between successive sighting points represent nightly movement distances, and the distance between the location of first capture (point 1) and location of final sighting (point 5) represents the displacement. Displacement ratio was calculated by dividing displacement by the sum of the nightly movement distances (a + b + c + d).

Because we had data on cephalothorax length and weight for different sub-sets of males, we tested for effects of both spider body weight and cephalothorax length (measured once at first capture) on mean nightly movement and mean nightly displacement using Pearson correlation analyses. Effects of environmental factors on nightly movement were analysed in two steps. First, we modelled a binomial variable (moved/not moved) to investigate the factors that affected whether spiders moved or stayed in the previous night’s location. Next, for spiders that had moved since the previous observation, we modelled a continuous (Gaussian) variable (nightly distance moved in m) to investigate the factors that affected how far spiders travelled while they were searching for mates.

The binomial and Gaussian models both contained the following environmental parameters as fixed effects: daily rainfall (mm), nightly minimum temperature (°C), daily maximum temperature (from the previous day; also, in °C), and plant community type. The models also included fixed effects of year, and order of days tracked for each individual (each day tracked for each spider was given a number starting at 1 on day 1 and increasing in numerical order). Both models included individual identification and a code representing each day when tracking occurred as random effects. The correlation between minimum daily temperature and maximum daily temperature was weak (R = 0.16), presumably because there was little seasonal change in average temperature during the time that spiders were tracked. We therefore included both the minimum and maximum daily temperature as fixed effects in our models. The plant community type (PCT) data obtained from NSW State Vegetation Type Map describes three broad habitat types present in Lane Cove National Park. These models were fitted using the TMB package (Kristensen, 2016).

For both the binomial and Gaussian response variables, we carried out model selection based on Akaike’s information criterion corrected for small sample size (AIC*_c_*) using the MuMIn package (Barton, 2022). We compared models that included all combinations of fixed effects (without interactions) and intercept-only models (Kristensen, 2016). All models included the two random effects, and diagnostics were assessed visually. Models with ΔAIC*_c_* < 2 relative to the best-supported model (i.e., the model with the lowest AIC*_c_* value) were considered to have some support, whereas models with ΔAIC*_c_* > 2 were considered to have very low support.

## Results

A total of 19 adult male *A. robustus* individuals were tracked between February and May of the years 2020, 2021 and 2022. Males used in this study were collected at night, typically as they were actively wandering along a trail or over the forest floor in search of a mate. After transmitter attachment and spider release, the males were tracked during the day. No males were sighted out and actively walking during the day; they were always found either inside a temporary shelter (see below) or, on 10 occasions, inside a burrow that was confirmed (in one case, described below) or inferred to contain a female. All burrows visited by males had characteristic *A. robustus* burrow features such as external thick silk sheeting, forming one or more funnels (entrances), multiple fine trip lines radiating out of each funnel up to ∼50 cm. Based on multiple features, such as absence of debris and the presence of white and reflective silk, these burrows could be deemed as active (Creak et al, in prep). We chose not to excavate or disturb these burrows to confirm female presence as this may have caused the female within to abandon the burrow, potentially impacting the movement of males nearby.

Male M8.22 was initially found and collected at night as he was approaching the burrow of a female. Once the transmitter was attached and he was released at the same location one week later, he immediately began courting by approaching the burrow and performing a series of leg movements. These movements resembled the behaviours of *A. robustus* males observed courting in the laboratory, especially ‘short leg contractions’ and ‘legs II quivering’ with the tarsi of legs I & II on the substrate/silk of the burrow (Frank et al., 2023). The female then responded by emerging from the burrow and performing several tapping movements, briefly touching the cephalothorax of the tagged male several times with her first or second pair of legs. As the male entered the burrow, a series of ‘body vibrations’ were performed where all legs briefly contracted and relaxed in a repeated pattern, causing the body to be lowered and raised several times while the tarsi remained in place.

### Burrow Visitation

In some cases, multiple males were tracked to the same burrow inferred to contain a female: males M2.22 and M6.22 were both tracked 19 days apart to one burrow, while males M11.21 and M12.21 were both tracked 15 days apart to the immediate vicinity of another burrow. There was also clear overlap in the areas these males were active (Figure 3), potentially indicating the presence of female burrows. No sexual cannibalism was observed, and all males tracked to burrows re-emerged and left the area to resume mate searching. Male M7.21 was tracked to two separate burrows seven days apart, and male M9.21 was also tracked to two separate burrows, eleven days apart. Several other males (n = 8) were each tracked to a single burrow. Time spent inside the burrow and in its immediate vicinity varied between one and five days.

**Figure 3:**
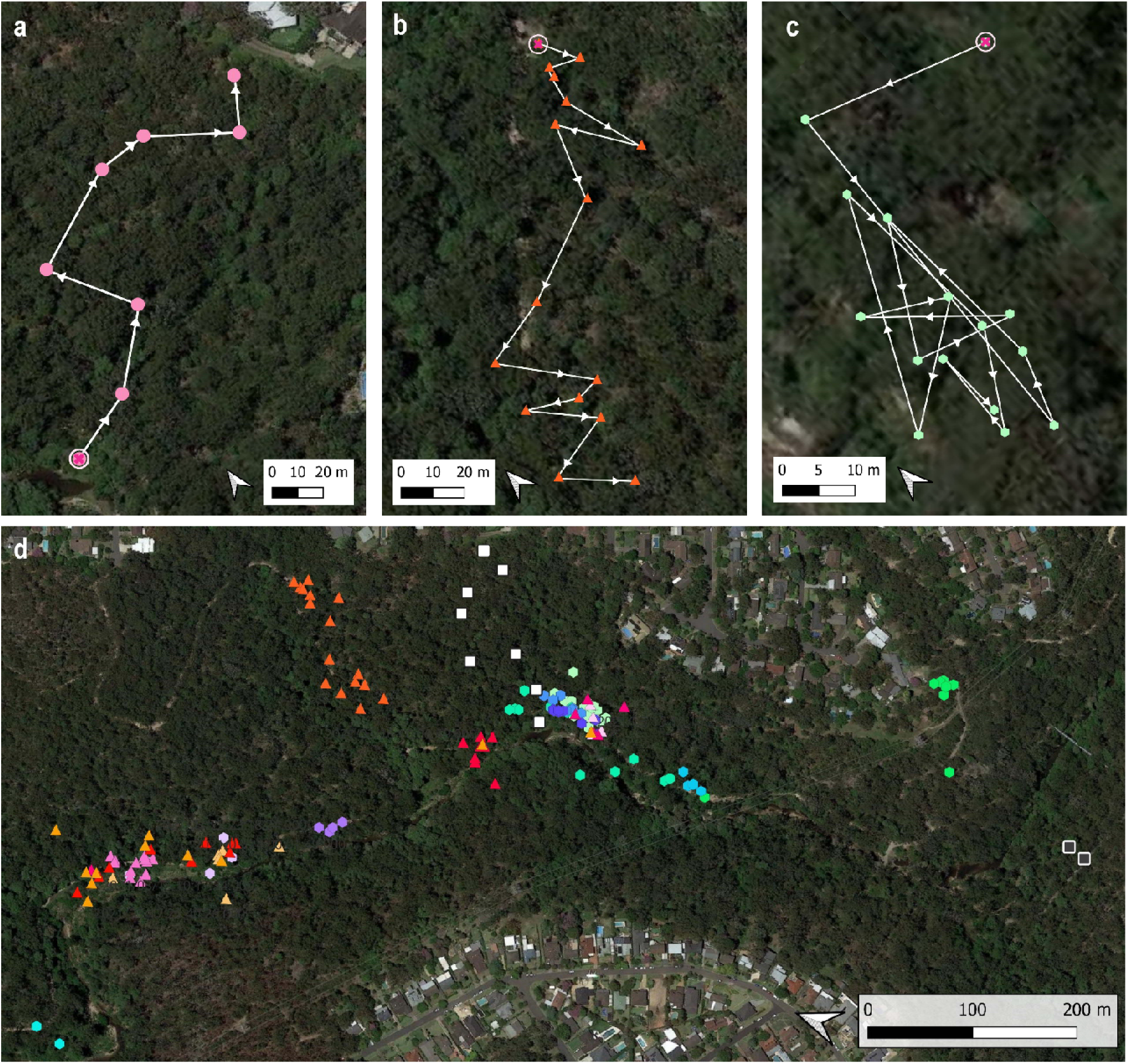
**a-c:** Examples of individual males’ movement patterns, with points showing GPS coordinates, arrows showing the direction of movement, and circled point showing the first point recorded (a; male M2.20. b; male M8.21. c; male M8.22). **d**; Map showing GPS location points for all males tracked during this study. Due to GPS error, several points were removed or manually corrected. Squares = males tagged in 2020, triangles = males tagged in 2021, hexagons = males tagged in 2022.

Two of the tagged spiders were killed by predators, and the transmitter was retrieved to confirm predation as the cause of death. The signal of the transmitter attached to M1.20 was detected from inside the stomach of a broad tailed gecko (*Phyllurus platurus*), indicating that the spider had been consumed (Amarasekara et al., in prep). The transmitter from M1.22 was found under dense leaflitter where, upon inspection, parts of the male’s cephalothorax were found still attached to the transmitter, indicating that the individual had been eaten and the transmitter discarded afterwards. The predator could not be identified in this case.

### Male spider movement patterns

The average nightly distance moved by male *A. robustus* was 11.56 meters per night. The longest distance covered by one individual (male M9.21) over a single night was 60.3 meters, while the shortest (male M11.21) was 0.4 meters. All spiders had a staggered type of nightly movement where nights without movement (when spiders were located on consecutive days inside a temporary shelter or burrow) alternated with nights when spiders moved to new locations. The probability of nightly movement was an average of 56.3%. On the occasions where males did not move and were not in the burrow of a female conspecific, they were often found in deep crevices against rocks or trees where they had created a rudimentary shelter consisting of a cocoon-like structure made from silk (see below).

Most individuals moved in a somewhat linear fashion, gradually moving away from their initial point of capture at an average rate of 4.01 m per night (Figure 3a and 3b). The mean displacement ratio was 0.69. However, one individual (male M8.22) moved in a circular pattern and ultimately ended up close to where he was first located (Figure 3c). We found that individual male weight was negatively correlated with mean displacement (r = −0.89, p = 0.04, Figure 4b) and (near-significantly) with the mean nightly distance moved (r = −0.82, p = 0.09, Figure 4a). However, there was no correlation between cephalothorax width and mean nightly distance moved (r = −0.07, p = 0.8 Figure 4c) or mean displacement (r = 0.22, p = 0.83 Figure 4d).

**Figure 4:**
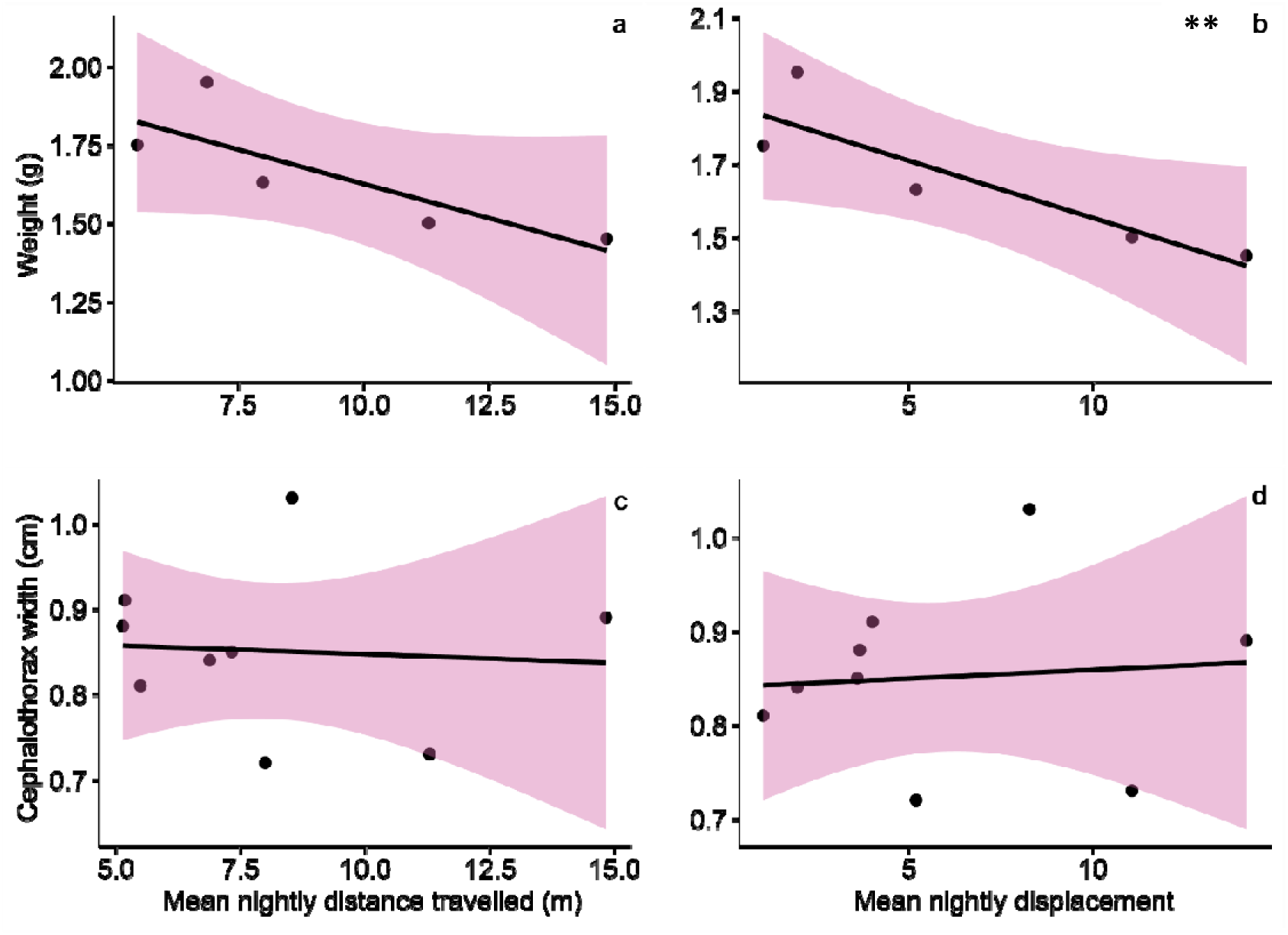
Pearsons correlations between mean displacement and weight (a), total distance travelled and weight (b), mean displacement and cephalothorax width (c), and total distance travelled and cephalothorax width (d). ** indicates a significant correlation. Points represent individual males, trend-lines are linear regressions, and shaded areas represent 95% confidence limits.

### Temporaculum structure and use

Tagged males located during the day, when not inside a female’s burrow, were typically found inside a temporary constructed retreat. We propose the term “temporaculum” for these temporary structures that serve as retreats between bouts of mate-searching (Figure 5). Whereas a hibernaculum is a structure in which an animal hibernates for long periods of time and/or frequently returns too, the temporaculum is not returned to by the individual, provides less protection, and requires less resource expenditure to create. *A. robustus* temporacula consisted of a fine layer of silk, often binding two or three leaves together, and were usually under the dense pockets of leaf litter found in rock crevices. Silk was also found attached to soil and rock potentially providing some anchoring and stability. Males spent between 1 and 6 (mean = 1.9) consecutive days in their temporacula before moving on, with the exception of M9.21 who spent 9 consecutive days in the same location.

**Figure 5.**
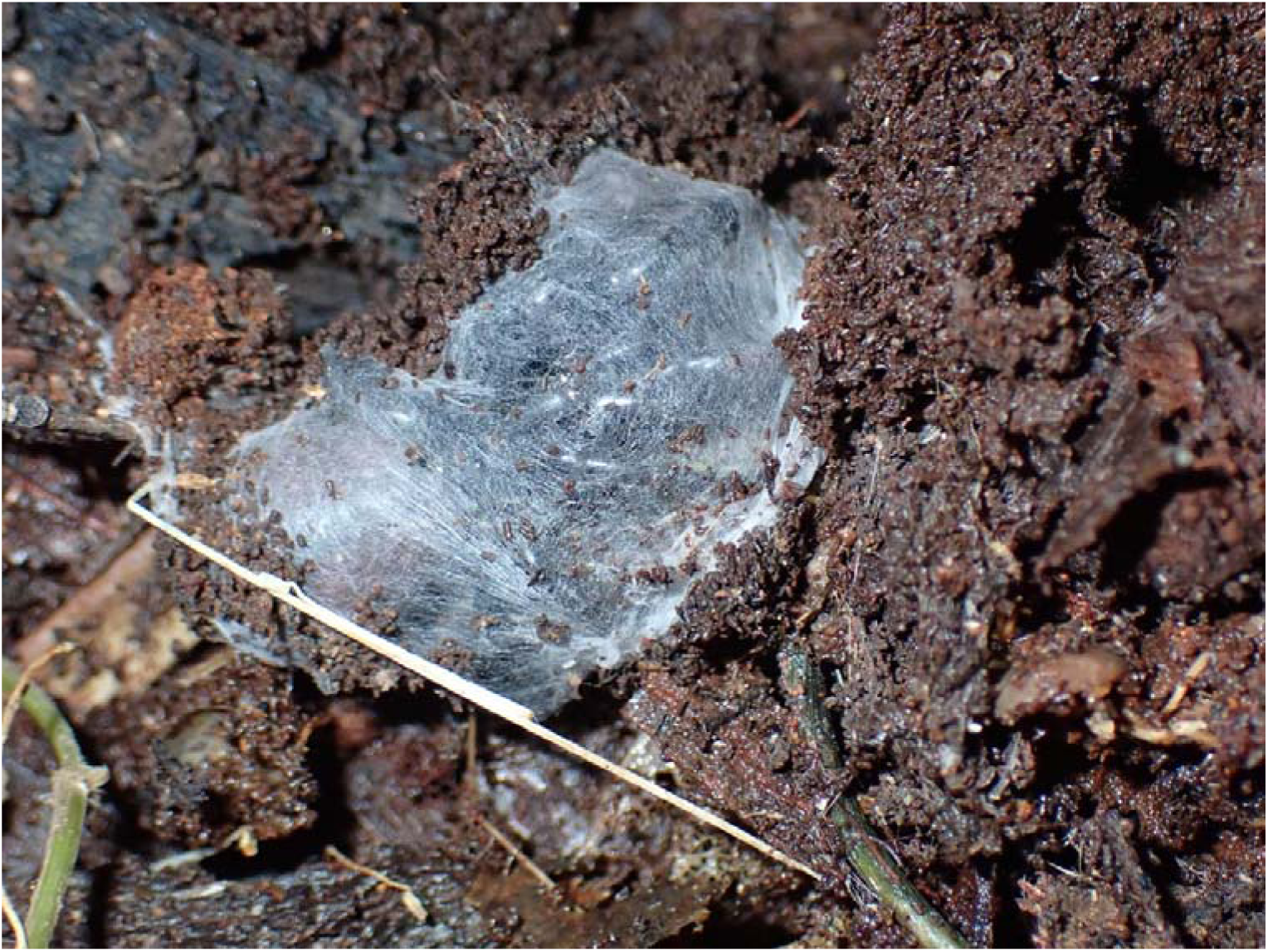
Photograph of a temporaculum of Atrax sp., with the male still enclosed inside it, found under a log in Bermagui NSW. The temporacula of A. robustus found in this study had a very similar structure to this.

### Effects of weather and habitat on male movement

The top model of the likelihood of movement during the night included maximum temperature (from the previous day), minimum temperature (overnight), and rainfall amount (Table 2). Rain reduced the likelihood of males moving, and a lack of movement was more evident during periods of heavier rain (Figure 6a). Spiders were more likely to move when the minimum daily temperature was high, and maximum daily temperature was low (Figure 7c, d). The only other factor included in supported models (i.e., models with ΔAIC*_c_* < 2) was order of day moved, which was included in one model with ΔAIC*_c_* = 1.14 (Table 2): the likelihood of moving increased with the number of days that individual males had been tracked (Figure 8a, Table 2). We found no support for an effect of plant community type (PCT) on the likelihood of an individual moving (ΔAIC*_c_* > 2 for models that include PCT, Table 2; Figure 8b).

**Figure 6.**
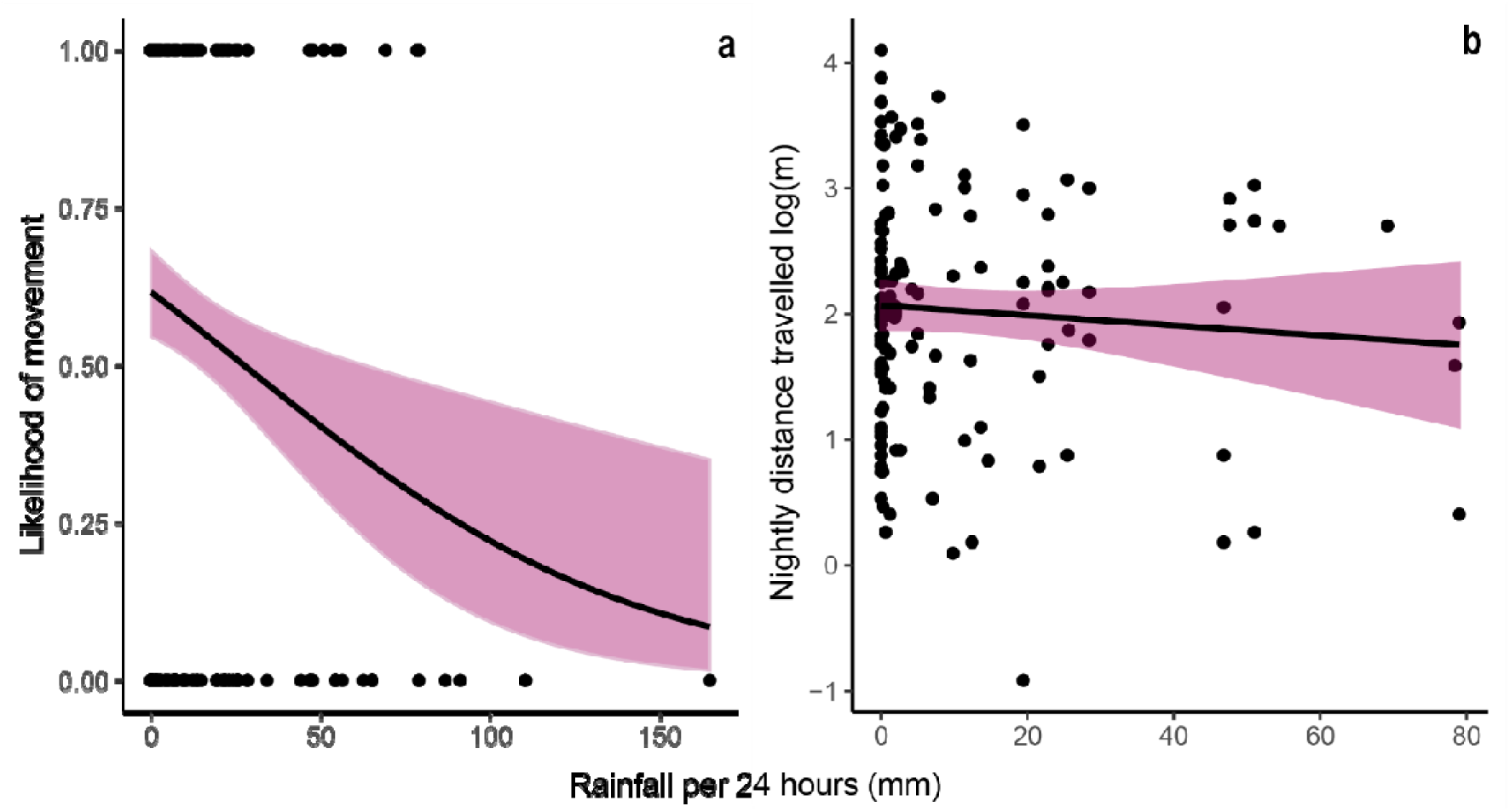
A Likelihood of movement based on rainfall per 24 hours. B Nightly distance travelled in meters (log-transformed)

**Figure 7.**
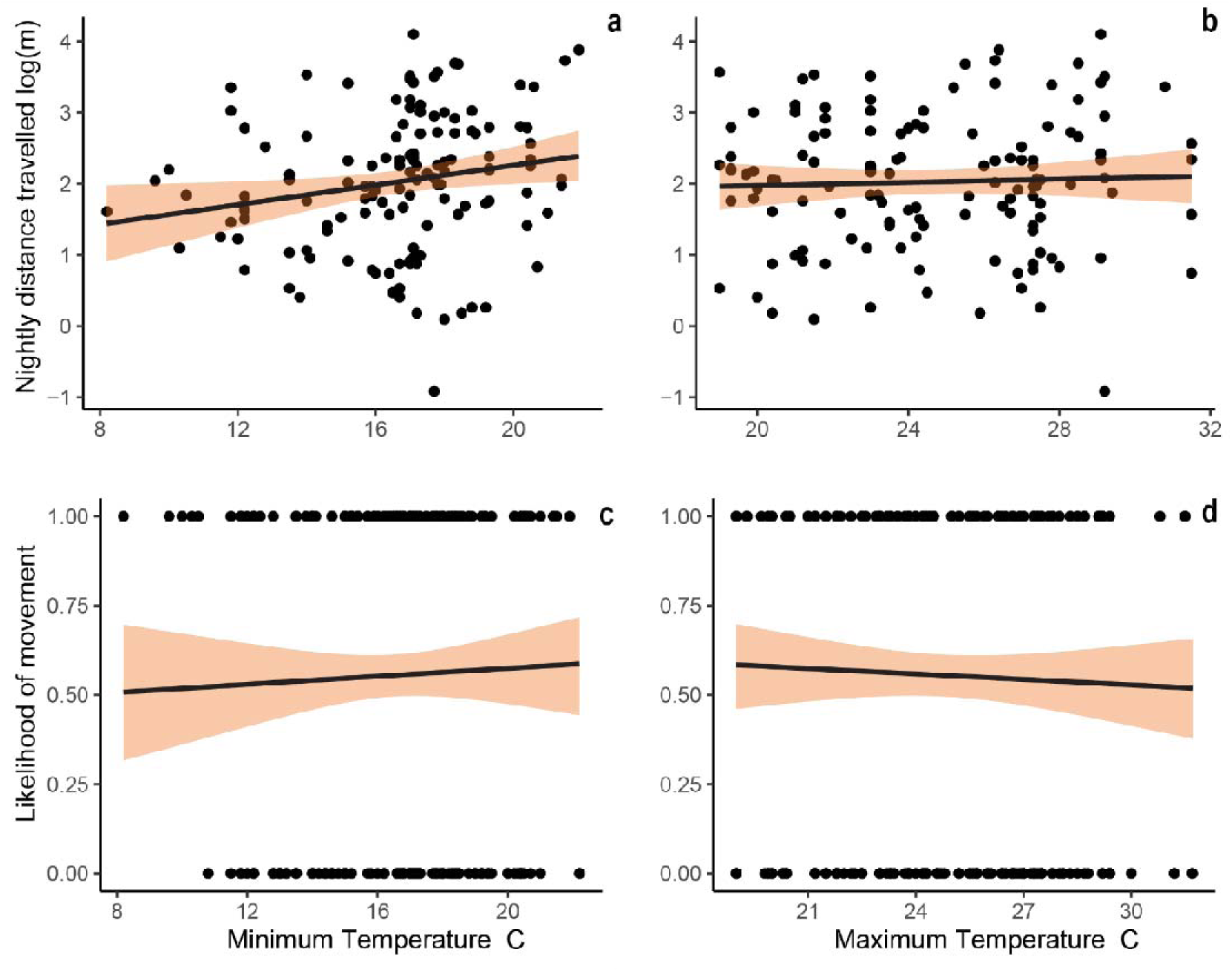
A) Nightly distance travelled and minimum temperature. B) Nightly distance travelled and Maximum temperature. C)Likelihood of movement and minimum temperature. D) Likelihood of movement and Maximum temperature. Trend-lines are linear regressions, and shaded areas represent 95% confidence limits.

**Figure 8.**
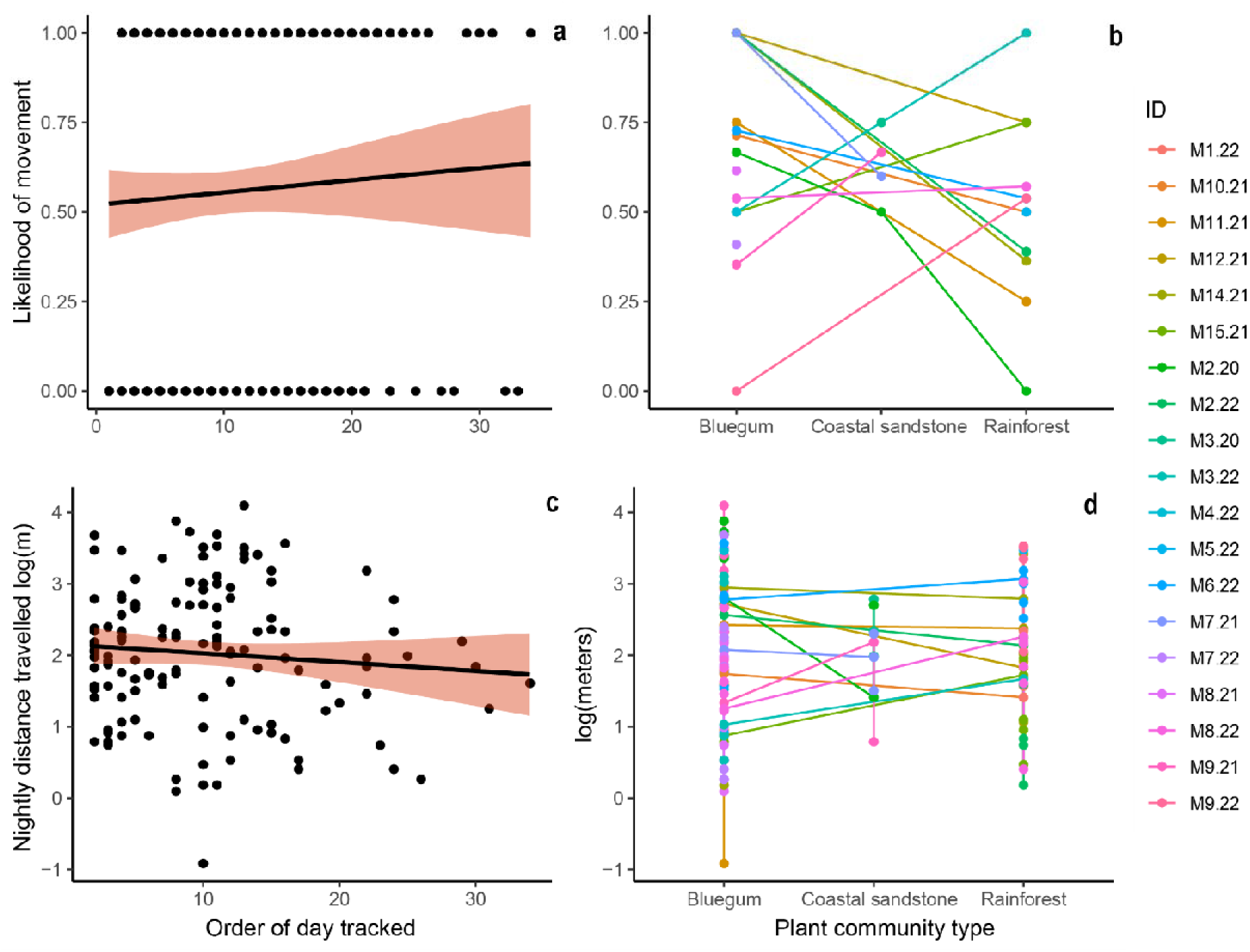
A) Likelihood of movement and the order of the days each individual was tracked. B) Nightly distance travelled in log transformed meters and the three plant community types, Bluegum, Coastal sandstone and Rainforest. Points represent individual nightly values. In panels a and c, trend-lines are linear regressions and shaded areas are 95% confidence limits. In panels b and d, points and lines are coloured by individual male ID.

**Figure 9.**
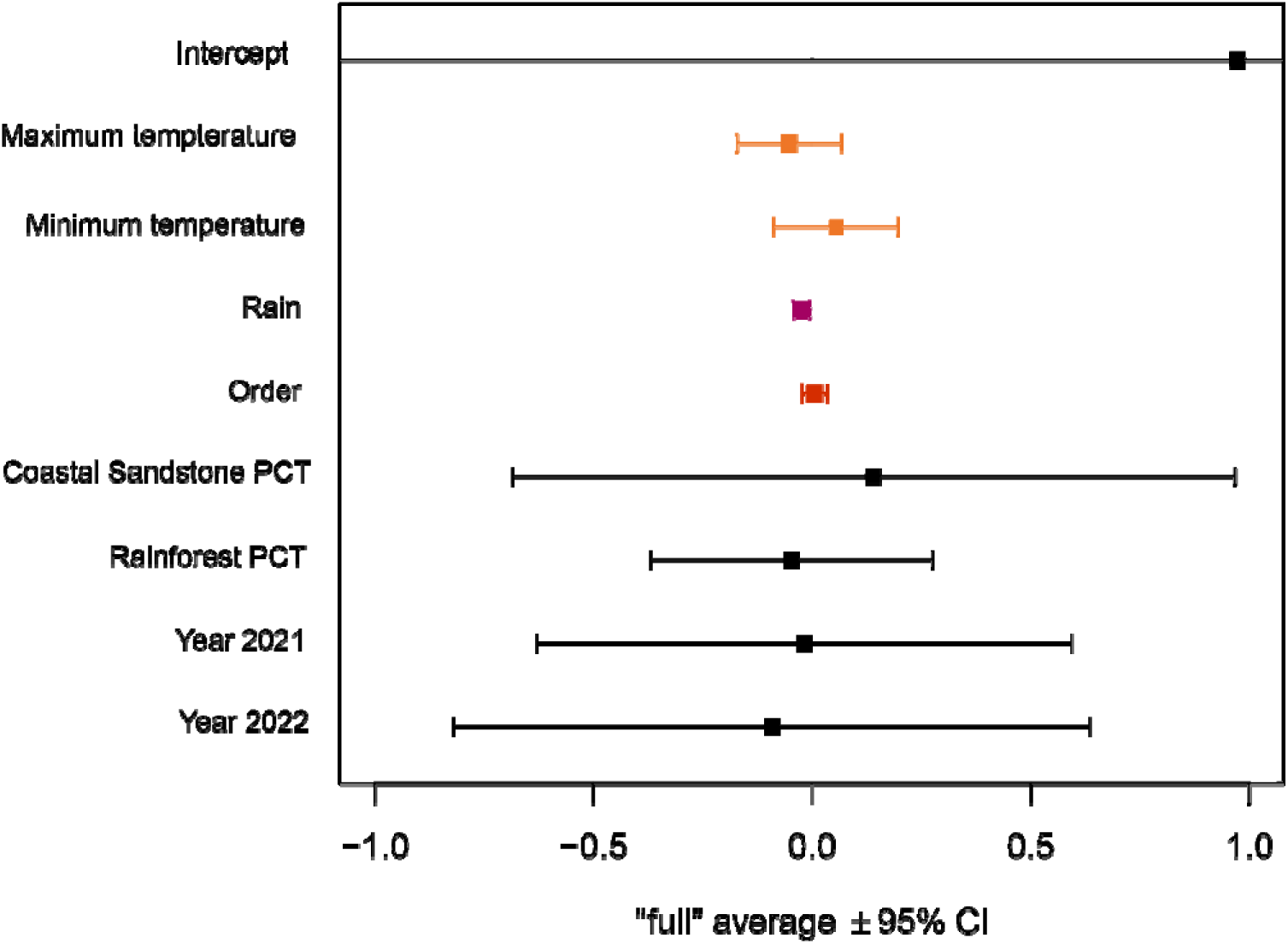
Model averaged coefficients for likelihood of movement

**Table 2:** Best-supported models examining effect of environmental parameters on the likelihood of movement. All subsets of the fixed main effects were compared. Only models with support (ΔAICc <2) are shown, as well as the null (intercept only) model.

| MODEL | DF | LOG LIKELIHOOD | AICc | $\Delta AICc$ | WEIGHT |
| --- | --- | --- | --- | --- | --- |
| MOVED ~ MAXIMUM TEMP + MINIMUM TEMP + RAIN | 6 | -178.816 | 369.9 | 0 | 0.14 |
| MOVED ~ RAIN | 4 | -181.292 | 370.7 | 0.79 | 0.09 |
| MOVED ~ MAXIMUM TEMP + RAIN | 5 | -180.400 | 371.0 | 1.08 | 0.08 |
| MOVED ~ MAXIMUM TEMP + MINIMUM TEMP + ORDER + RAIN | 7 | -178.331 | 371.1 | 1.14 | 0.08 |
| MOVED ~ MINIMUM TEMP + RAIN | 5 | -180.513 | 371.2 | 1.30 | 0.07 |
| INTERCEPT ONLY | 3 | -185.746 | 377.6 | 7.63 | 0.00 |

The top model of nightly distance travelled included only the minimum temperature: spiders covered less distance on cold nights (Table 3; Figure 7a). Other supported models (i.e., models with ΔAIC*_c_*< 2) also included rain, year, PCT and order of days moved (Table 3). Rainfall amount was included in one of the best-supported models (ΔAIC*_c_*= 0.13): the nightly distance travelled was negatively affected by amount of rainfall (Figure 6b). PCT was also included in one of the best-supported models (ΔAIC*_c_* = 0.07, Table 3), with the greatest distance travelled in the Rainforest PCT (Figure 8d; Figure 10). We also found some evidence that the nightly distance travelled decreased with the number of days tracked (ΔAIC*_c_* = 1.8 relative to the best model, Figure 8c; Table 3). Nightly distance travelled was not affected by the daily maximum temperature (ΔAIC*_c_* > 2 for models that include maximum temperature; Table 3; Figure 7b).

**Figure 10.**
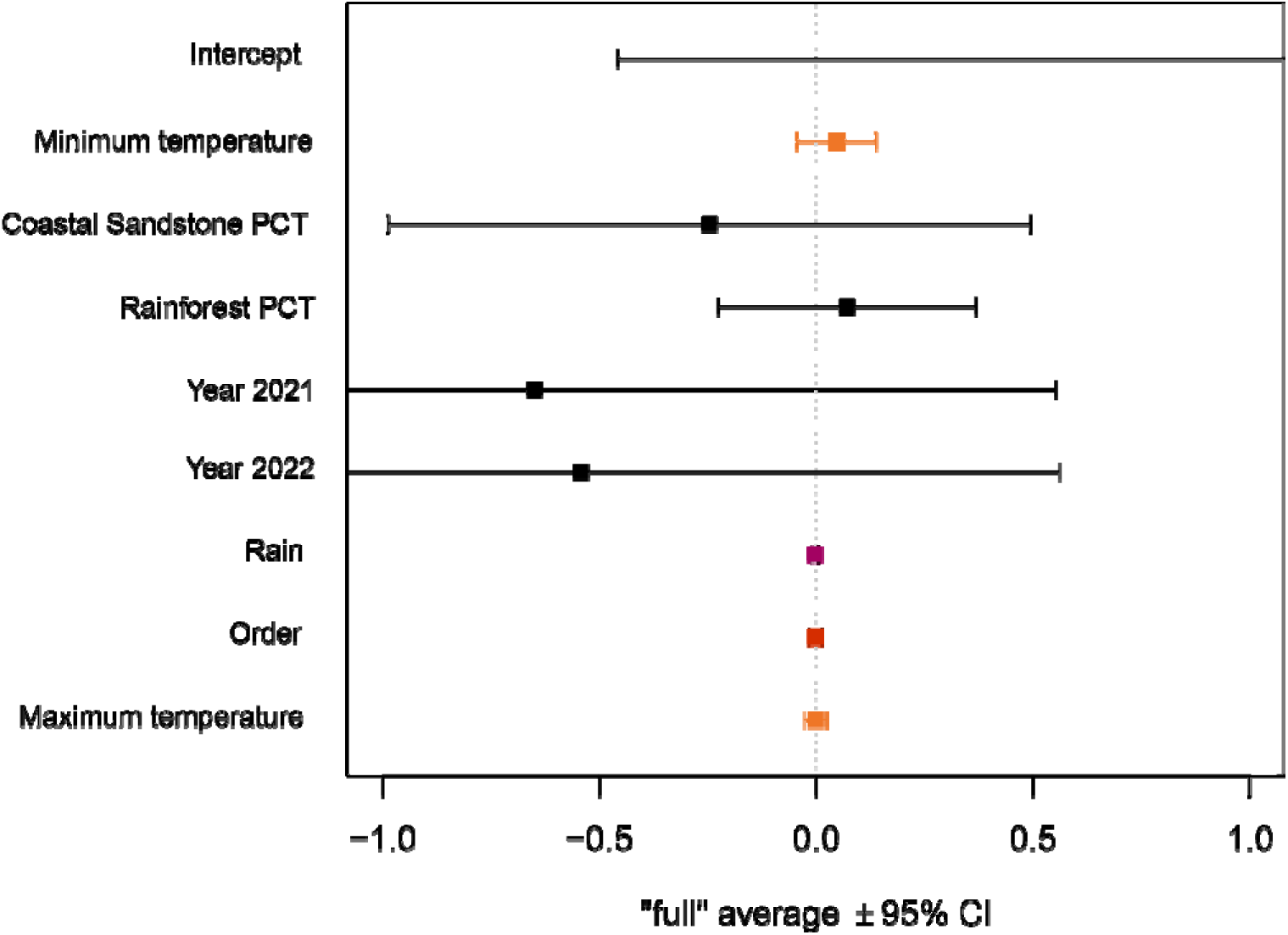
Model averaged coefficients for Nightly distance travelled

**Table 3:**
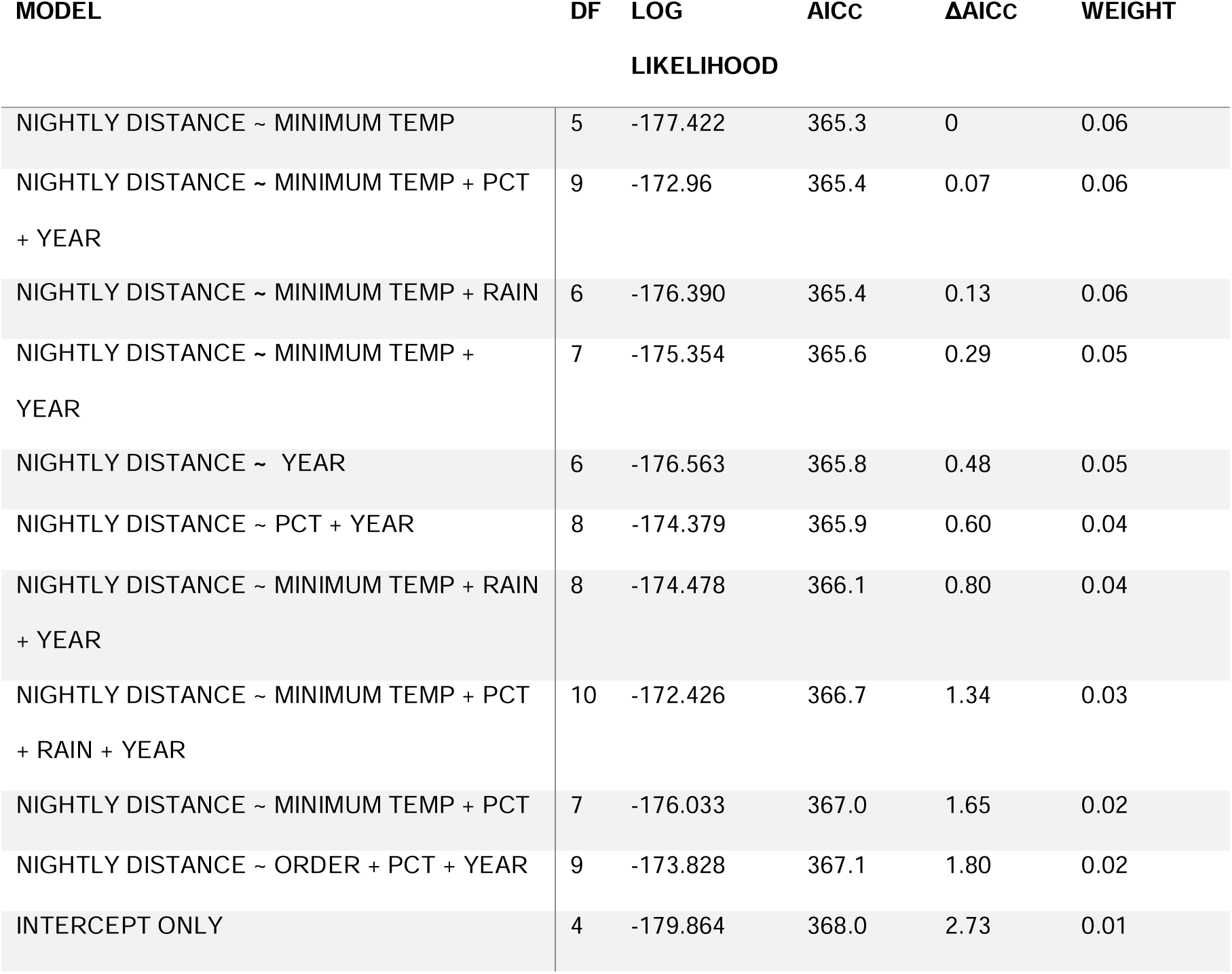
Best-supported models examining effect of environmental parameters on the nightly distance travelled by individuals. All subsets of the fixed main effects were compared. Only models with support (ΔAICc <2) are shown, as well as the null (intercept only) model.

## Discussion

Very little is known about the movement patterns spiders during mate-searching in their natural habitat. The goal of this study was to help fill that gap by investigating the active mate searching behaviour of the Sydney funnel-web spider *Atrax robustus* and testing the prediction that mate searching behaviours minimize risk while still enabling males to locate mates. Our study contributes data on the behaviours and strategies used by males during the phase between the commencement of mate-searching and the location of a mate, as well as evidence of how this movement is affected by environmental conditions. We found that males only searched for mates at night and remained sheltered during the day in temporary shelters constructed from silk. *A. robustus* males were less likely to move during rainy periods and nightly distance moved also decreased with heavier rainfall. Males were more likely to move and moved greater distances during warmer nights, and the likelihood of movement was also greater on nights that followed cool days. These findings suggest that the behaviour of the mate-searching sex appears to minimize risk during periods of active mate-searching.

Burrows visited by *A. robustus* males were inferred to contain adult females (confirmed in one case by direct observation), and such burrows sometimes received sequential visits from multiple males. This suggests that *A. robustus* females can mate with multiple males, generating an opportunity for sperm competition and thus favouring mate guarding behaviours in males. Indeed, some males in this study were found for up to six days in and around burrows inferred to contain adult females, suggesting a prolonged mate guarding behaviour. Mate guarding has only been observed in one other species of mygalomorph, *Mecicobothrium thorelli* (Mecicobothriidae), where males only guarded the female burrow for approximately 30 minutes after copulation (Ferretti et al., 2013). More research is therefore needed to determine whether the multiple mating and mate guarding behaviours observed in *A. robustus* are typical of other mygalomorph spiders.

Temporacula (temporary shelters built from silk and leaf-litter, where males often sheltered during the day) were created and used by male *A. robustus* every day that they were not found inside of a potential mate’s burrow. With *A. robustus* presenting a staggered pattern in movement frequency, temporary retreats served as a protective shelter. The construction of temporary retreats by searching male spiders has not been recorded in other species of mygalomorphs. However, in one species of tarantula (family Theraphosidae), *Aphonopelma anax*, males who abandoned their burrows upon maturation sheltered in a variety of existing structures, such as abandoned mammal burrows and, similar to *A. robustus*, did not return to the same daytime shelter (Shillington, 2002). Shillington (2002) hypothesized that males were using these temporary shelters to conserve energy, since males that utilized the temporary shelters performed better during experimental trials. To our knowledge, the use of temporary shelters by searching males has not been reported in other spiders. However, in many species of jumping spider (Salticidae), individuals have been recorded creating and utilizing hibernacula (Hoefler & Jakob, 2006).

The three plant community types that *A. robustus* males were observed traversing in this study initially appeared to influence the nightly distance moved, since males appeared to favour moving in the rainforest habitat. While we expected habitat types and their varying complexity to influence the mate searching behaviours of males, the PCTs obtained from NSW State Vegetation Type Maps (State Government of NSW and NSW Department of Climate Change, Energy, the Environment and Water, 2022) depict a large-scale resolution that did not capture smaller plant communities such as dense clumps of pteridophytes (ferns and relatives) or areas predominantly covered with *Allocasuarina* spp. In addition, the area in which this study was conducted is identified as primarily bluegum PCT, while the other PCTs cover relatively small areas of the study. Male *A. robustus* can disperse tens of meters per night and although they may stay in one broad PCT for several days or weeks, they are likely to encounter a mosaic of small-scale habitat types each night. Preferred habitat type for burrow construction is documented in several species of mygalomorphs (Beavis et al., 2011; M’rabet et al., 2007) and is vital for conservation efforts, antivenom collection, and education. Our findings suggest that although there may be a relationship between mate searching behaviour and PCT, further investigation with a finer resolution of habitat type is needed.

Tagged male *A. robustus* individuals were half as likely to move during periods of heavy rain as on dry nights, and in cases where movement was observed, shorter distances were travelled on rainy nights. Although there is evidence to suggest that female *A. robustus* may be more active after heavy rainfall (Bradley, 1993), our results show that male activity decreases during or after heavy rain. There are several unpublished reports of *A. robustus* surviving under water in pools for at least 30 hours (Australian museum, 2022). However, males may still be vulnerable to a variety of other biotic and abiotic factors during heavy rainfall, such as fast-moving floodwater and the debris often accompanying it. Our results suggest that *A. robustus* males avoid mate-searching during such risky weather conditions.

*A. robustus* males exhibited a zig-zag pattern of movement that tended to result in gradual displacement away from their point of initial capture. A zig-zag movement pattern has been observed in a multitude of mate searching invertebrates in response to volatile pheromone and is believed to optimize the chances of intercepting the source (Kanzaki, 1998). Since a two-dimensional environment makes it easier to follow directional pheromonal cues, ground dwelling spiders may be expected to move in a more linear direction in comparison to aerial web builders such as orb weavers. This prediction could be tested by comparing movement of mate-searching males in two- vs three-dimensional environments. The touch pheromones typically found on silk or on the cuticle of the individual have been found to become inactive after being exposed to water, while volatile pheromones quickly dissipate in open bushland (Dondale & Hegdekar, 1973; Gaskett, 2007). In contrast, *Eupalaestrus weijenberghi*, a tarantula found in meadows in Uruguay, was found to have an extremely persistent pheromonal signal that lasted up to 55 days in natural stormy conditions, and *E. weijenberghi* males placed in terraria with rain-exposed female burrows still exhibited courtship behaviour (Costa et al., 2015). This zig-zag movement, in conjunction with significantly reduced likelihood of movement during rainy periods, is suggestive of a non-water-resistant pheromone being emitted by female *A. robustus.* Males may remain stationary until females produce fresh pheromone trails, thus reducing energy expenditure and risk.

Tagged males were more likely to move and moved greater distances on warm nights, and were also more likely to move on nights that followed cool days. During summer (December – February) in the Sydney region, temperatures can reach >40°C during the day and drop to ∼15°C at night. Despite large fluctuations throughout the day, temperatures at night are more consistent. *A. robustus* males permanently leaving their burrows to search for a mate are thus exposed to the risk of heat stress during the day. This could explain why nocturnal mate searching is preferred by *A. robustus* males. In addition, the risk of predation might be reduced by mate searching at night, especially if birds are important predators of *A. robustus*. Although one of the tagged males was eaten by a nocturnal gecko, the main predators of *A. robustus* are still unknown. Mate searching during a specific time of day can also be seen in other species of mygalomorphs that encounter fluctuating temperatures as they often have preferred temperature ranges (Alfaro et al., 2013). Temperature also plays a role in female mate receptivity and reproductive success in other species of mygalomorph (Veloso et al., 2012). In recent studies there has been a consistent trend showing spider vulnerability to high temperatures due to water loss, particularly in mygalomorphs (Mason et al., 2013). Changes in sexual behaviour in response to fluctuating temperatures have also been observed in other species of spiders, where courtship behaviours have declined in quality with rising temperatures (Jiao et al., 2009). Travelling further distances within a particular temperature range suggests that mate searching behaviour may be restricted due to thermoregulatory requirements and provides further evidence that males search for mates when risks are low.

During our study we observed an extreme early emergence of a mate searching male *A. robustus*, indicating potential effects of climate change on this species. During June of 2023, typically the coldest month in NSW, a sexually mature male *A. robustus* was collected by a volunteer in Hornsby Heights NSW. This was the first time that an *A. robustus* male had been observed mate-searching during the winter. This early emergence may have been induced by the record-high winter temperature in NSW, 2.33°C above the historical average (Australian Government, Bureau of Meteorology, 2023).

The mate searching behaviour of male *A. robustus* is of particular interest given the lethal and complex nature of their venom (Escoubas et al., 2006; Pineda et al., 2020). The distribution of *A. robustus* spans over some of the most densely populated areas of Sydney and the collection of wild adult males is essential to the production of a lifesaving antivenom. Understanding how environmental conditions affect the mate searching activity of male *A. robustus* could result in searches that are optimized for collecting males of this species with as little ecological disturbance as possible. Although there have been some research on the behaviour of mygalomorph spiders, nearly all of these studies have been conducted in laboratory environments. More research is needed to understand how their behaviours function and vary in a natural and changing habitat. In particular, there is very little information available across all animals on behaviours and strategies employed during the active mate searching period. Research on *A. robustus* will help to fill the gap in knowledge, but further studies are needed.

Today, very little is known about the behaviour of *A. robustus* in its natural habitat, since recent studies have only investigated individuals under laboratory conditions to assess behaviours such as defensiveness, activity, aggression (Duran et al., 2022, 2023; Hernandez Duran et al., 2023) and courtship (Frank et al. 2023). Our study provides preliminary insights into the behavioural ecology of *A. robustus* males in a natural setting, and reveals several novel behaviours and strategies used by males of this species. In particular, we observed a novel form of prolonged mate guarding, unknown in other species of mygalomorph spider. We also discovered that *A. robustus* males construct temporary shelters (temporacula). Our tracking data suggest that male *A. robustus* individuals locate mates by following volatile pheromone cues, like many other arthropods. Finally, we provide several lines of evidence suggesting that males minimize risk during their active mate searching period: risk minimization is suggested by the observations that males construct temporary shelters (temporacula), and are more likely to search for females, and/or move further distances, in preferred weather conditions (warm, dry nights following cool days).

## Acknowledgements

We acknowledge the traditional custodians of the land on which this research was conducted, the Dharug peoples, and that traditional knowledge and stories of this species amongst many others may have been lost. We thank the numerous volunteers that assisted with fieldwork, Melissa Abdallah, Sophie Ren, Daniel Creak, Joshua Creak & James Gallagher. We also thank Braxton Jones for notes and insights. We would like to thank UNSW Stats Central and Dr Benjamin Walker for assistance with statistical analysis, and National Parks and Wildlife Services for supporting our work in Lane Cove national park. This research was funded by National Geographic Society (project number GR-000046126) and Australian Geographic Society (103961107).

